# Septin inhibitors exert opposing effects on distinct higher-order septin structures in melanoma cells

**DOI:** 10.64898/2026.09.08.750271

**Authors:** Maggie M. Gravano, Leah M. Farmer, Nathaniel Richmond, Ishaan Mehta, Anushka Deshmukh, Alecxander J. Lewis, Andrew D. Weems

## Abstract

Septins are cytoskeletal GTPases that assemble into diverse higher-order structures with context-dependent mechanical and signaling functions. Recent work identified a septin-dependent pro-survival pathway in metastatic cancer cells, termed *bleb signaling*, in which plasma membrane blebbing promotes the formation of septin signaling hubs that support survival during cellular detachment and dissemination. Disrupting these hubs may offer a novel therapeutic strategy targeting disseminated cancer cells. Here, we compare the effects of three septin-targeting compounds, forchlorfenuron (FCF), UR214-9, and REM127, on septin organization and viability in melanoma cells using confocal microscopy and quantitative computer vision analysis. FCF and UR214-9 both effectively destabilized septin hubs, while REM127 had no appreciable effect. FCF also reproduced the context-specific viability reduction previously seen in detached melanoma cells, while UR214-9 caused broad off-target toxicity and cell death in all growth conditions. As with hub disruption, REM127 produced no effect on viability. Surprisingly, though none of the compounds caused an increase in higher-order septin structures during detachment, all three increased end-on-end polymerization of filamentous septins at stress fibers when grown adhered to substrate. These findings demonstrate that septin-targeting compounds exert divergent effects on different classes of septin higher-order structures within the same cell type, promoting filamentous assembly while disrupting or failing to promote membrane-associated signaling hubs. This highlights not only the structural distinctiveness of such classes, but that pharmacological targeting can produce opposite effects on them. Moreover, this study emphasizes the need for next-generation septin inhibitors capable of selectively targeting disease-relevant septin assemblies.

## Introduction

Septins are GTP-binding cytoskeleton proteins that can assemble into filaments and other higher-order structures^1^. They exist predominantly as rod-shaped hexameric or octameric palindromic heterooligomers. Interactions between the outer subunits of these heterooligomers allow for end-on-end inter-oligomer polymerization to form higher-order septin structures^2–4^. In different cellular contexts septins can fill both mechanical and signaling roles. They can bind to and polymerize along both F-actin and microtubules^5–8^, contributing to organizing the cytoskeleton^9^, anchoring it to the plasma membrane^7^, and buffering against mechanical forces^10,11^. They also serve as signaling scaffolds capable of regulating a wide variety of signaling pathways, by binding with and modulating the local concentrations of signaling factors and their various effectors^12^. Interestingly, septins are also the only eukaryotic proteins known to be capable of sensing micron-scale membrane curvature, consistently accumulating on the plasma membrane at such sites^13,14^. Septin involvement in numerous cellular processes, including cell division^15,16^, polarization^17^, host–pathogen defense mechanisms^18,19^, primary cilia formation^20,21^, and cell migration^22^, has increased interest in understanding septin dynamics. Importantly, septin function has also been linked to various aspects of cancer disease progression, though the cell biological mechanisms underpinning these septin-cancer associations are not fully understood^23–25^.

Recent work by our group has revealed a septin-mediated cancer signaling phenomenon which might partially bridge this conceptual gap. We found that metastatic cancer cells often construct expansive septin structures at the plasma membrane, where they form signaling hubs capable of driving inappropriate cell survival in challenging microenvironments such as the stroma and sites of distant metastasis. This process is enabled by the amoeboid morphological program and the dynamic blebbing of the plasma membrane it brings about. Such blebs generate the specific scales of membrane curvature that attract septins, allowing sustained blebbing to serve as a morphological activation signal for the construction of septin hubs and the downstream signals they generate. For this reason, we named the novel signaling pathway “bleb signaling”^12^. Follow-up studies using human patient samples and mouse models of metastasis confirm that this phenomenon is employed by metastatic cells *in vivo*, with the pathway specifically becoming activated as cells detach from the main tumor mass at the invasive margin and staying active throughout the metastatic cascade^26^. Importantly, we found that pharmacological septin inhibition in our mouse model resulted in the specific killing of disseminated cancer in the stroma and at sites of distant metastasis, reducing metastatic burden and prolonging disease-free survival after tumor resection. These results highlight septins as a promising new therapeutic target in metastatic cancer.

The most well characterized septin inhibitor is forchlorfenuron (FCF). First identified as a synthetic cytokinin that enhances plant growth^27^, FCF has been used in agriculture for decades. FCF alters septin localization and stabilizes septin polymers in fungal, nematode, arthropod, amphibian, and mammalian cells, including human cell lines^28–30^. FCF’s exact mechanism of action is unknown, but *in silico* simulations predict that it interacts with the septin nucleotide binding pocket^31^. FCF selectively induces cell death in detached but not adherent melanoma cells by disrupting membrane septin hub formation^12^. It is not fully understood why FCF stabilizes higher-order septin assembly in some contexts but paradoxically disrupts membrane hubs, though this septin destabilizing effect has also been observed in other cellular contexts^17,24,32^.

FCF is not the sole small molecule inhibitor of septin function, with several groups publishing novel inhibitors in recent years. Two of the most promising septin inhibitor candidates have been UR214-9 from the UR214 series and REM127 from the Res19-T series. UR214-9 showed improved inhibition of septin function in cell lines at a reduced concentration compared to FCF^33^. However, later work has raised concerns of general cytotoxicity of UR214-9 in both cell lines and zebrafish embryos^34^. Similarly, REM127 restored septin filament integrity in Alzheimer’s disease models, normalizing calcium dysregulation^35^, but it was later withdrawn from a Phase 2a Alzheimer’s trial due to liver toxicity^36^. Decades of agricultural safety studies by the EPA in the US and the EFSA in the EU suggest FCF is relatively non-toxic, though its clinical use has never been tested by the FDA^37,38^.

The mechanism of UR214-9 has not been directly characterized, but as an FCF derivative, they are assumed to have a similar mechanism. Alternatively, REM127 has been proposed to act as a “molecular glue” between SEPT6 and SEPT7 within the septin oligomer^35^. Neither compound has yet been tested for effects on bleb-generated septin hub formation. To better understand how these septin-targeting compounds affect higher-order septin structures in a bleb signaling context, we evaluated UR214-9 and REM127 compared to FCF on septin hub formation in MV3 melanoma cells.

## Methods

### Cell culture

MV3 cells were a gift from Peter Friedl (MD Anderson Cancer Center, Houston, TX). Cells were cultured in DMEM (ThermoFisher, #11995073) supplemented with 10% fetal bovine serum (Peak Serum, # PS-FB2) and antibiotic-antimycotic (Gibco, #15240062) at 37 °C and 5% CO_2_ and grown on 10cm tissue culture dishes (VWR, #10861-594). All cell lines were tested monthly for mycoplasma contamination using the PCR Mycoplasm Detection Kit (ThermoFisher, #J66117.AMJ). Expression of SEPT6-eGFP in MV3 cells was achieved through lentivirus transduction as previously described^12^.

### Microscopy

All images were acquired using a Nikon Eclipse Ti2 microscope controlled by Micro-Manager software^39,40^. Cells were imaged using an Okolab stage-top CO_2_ incubator maintained at 5% CO_2_ and 37 °C. A 100× silicone oil immersion objective (Nikon, Silicone Immersion Oil 30cc MXA22179) was used for all experiments, except for the viability assay, which used a 10× air objective lens. The microscope was equipped with a Crest X-Light V3 spinning disk confocal head and a Photometrics Prime95B camera. SEPT6-eGFP was excited using a 488 nm laser, Propidium Iodide stain with a 555 nm laser, and Hoechst with a 405 nm laser. The FIJI ImageJ package^41,42^ was used to view and analyze Z-stack acquisitions and prepare maximum-intensity projection and slice images.

### Inhibitors

Drug compounds were reconstituted in Dimethyl sulfoxide (DMSO) [Sigma, #D1435-500ML], sonicated, and filter sterilized. Reconstitutions were stored at −80 °C and thawed the day of use. Concentrations used for each experiment were 50 uM Forchlorfenuron (FCF) [Sigma, #C2791], 25 uM UR214-9 [TCG Lifesciences], and 200nM REM127 [TCG Lifesciences]. Drug treatments added to cells were freshly prepared in phenol red-free DMEM containing 25 mM HEPES (ThermoFisher, #21063029) with 10% FBS and antibiotic–antimicotic (ThermoFisher, #15240062) and 0.1% DMSO. The solvent-only control medium was also prepared at 0.1% DMSO.

### Detached cell imaging and manual cell classification

SEPT6-eGFP MV3 cells were grown to 70–80% confluency in 10cm dishes, trypsinized for 3-5 minutes, and then seeded onto 60mm Ultra-Low Attachment dishes (Corning, #3261) at approximately 25,000 cells ml−1 based on Trypan blue counts (MP Biomedicals, #1691049). They were incubated in this detached state for two hours in a drug-treated medium or DMSO solvent-only control medium. Following incubation, the cell suspension was transferred to a 35mm poly-D-lysine coated glass-bottom dish (MatTek, #P35G-1.5-14-C), and all images were acquired within 30 minutes. Cells were visualized with the FIJI ImageJ package and scored according to a standardized protocol. Cells not fully in frame across all time points, cells making contact with other cells, cells beginning to show signs of adhesion to the dish, cells lacking membrane blebs, and cells that appeared dead or displayed grossly abnormal morphologies suggesting the activation of programmed cell death were all excluded from quantification. All cell classifications were performed blind to drug treatment conditions. For human-rated calls, cells were classified as bleb signaling active when fluorescent puncta corresponding to septin enrichment at the membrane were present, and as bleb signaling inactive when such puncta were absent.

### Computer vision analysis of detached cell images

We supplemented manual visual calls for bleb signaling by developing a computer vision pipeline to sample SEPT6-eGFP signal intensity at the cell membrane. This approach allowed us to extract 3D fluorescence data mapped to the cell surface, enabling a precise comparison of septin intensity, and relative enrichment between the membrane and the cytosol.

To sample SEPT6-eGFP intensity at the cell membrane, we generated 3D surface meshes from cell segmentations to serve as spatial references. Segmentations were produced via a multi-step pipeline. First, raw anisotropic images were upsampled in Z by a factor of 9.09 to yield isotropic 0.11 µm voxels matching the lateral pixel size. Next, training data was generated by extracting and manually segmenting 30 slices from each principle orthogonal view (XY, XZ, and YZ) in Napari^43^. Because axial cross-sections show a markedly different appearance from lateral ones, these annotations were used to train two custom Cellpose models^44–46^ initialized from the cyto3 pretrained models: one for XY slices, and a second trained on the pooled XZ and YZ annotations. Both were trained for 1000 epochs (learning rate 0.002, weight decay 1e-4, momentum 0.9, batch size 8) with a 90/10 training/validation split. The trained models were then applied to every slice of every volume along all three orthogonal planes (flow threshold 0.6, cell probability threshold −1, minimum object size 5 px, diameter taken from the trained model). Finally, the three orthogonal stacks of 2D labels were aggregated into a cohesive 3D instance segmentation using the indirect aggregation method of u-Segment3D^47^.

Surface meshes were extracted from each binary 3D segmentation by marching cubes on a Gaussian-smoothed mask (σ = 2.0 voxels, contour level 0.5). The resulting surfaces were isotropically remeshed (CGAL, minimum 10,000 elements) so that vertices were uniformly distributed over the cell surface. Meshes remained in voxel coordinates so that vertex positions match the isotropic image volume directly, avoiding any interpolation between mesh and image space.

Prior to intensity sampling, we applied conformalized Mean Curvature Flow (cMCF) to each mesh to derive a smoothed reference surface (S_ref) that removes fine protrusions while preserving overall cell shape. The Signed Distance Function (SDF) of the segmentation was computed and its gradient evaluated at each S_ref vertex to define the local inward sampling direction. The SDF approach has the added benefit of preventing sampling trajectories from crossing each other. To ensure reliable data, vertices were retained for analysis only if they satisfied two criteria: (i) they lay within the middle third of the cell in Z (axial centroid ± one-sixth of the cell height), excluding apical and basal surfaces where axial resolution is poorest; and (ii) the SDF gradient at the vertex was anti-aligned with the outward surface normal (cosine < −0.5),ensuring each ray traversed along the true inward direction rather than obliquely.

The reference surface was then superimposed onto the isotropic raw images, where SEPT6-eGFP intensity was sampled at each retained vertex along rays extending 3 µm inward and 3 µm outward, following the SDF gradient in steps of 0.5 voxels (0.055 µm). A cytosolic reference was established per cell from the pooled intensity samples of all retained rays falling 2.5-3 µm inward of the surface. Within this 0.5 µm range each ray contributed 9 intensity samples, which were pooled across thousands of retained vertices. Because this sampling region may contain cytosolic volumes, low intensity nuclear volumes, and occasional high intensity intracellular structures, we took the 75th-85th percentile band of the pooled distribution (which we determined through trial and error to yield the most accurate measurement of diffuse cytosolic signal in all experimental groups and replicates) and used its mean (µ_cyto) and standard deviation (σ_cyto) as the reference.

Membrane enrichment was then called independently for each ray. Within a surface window spanning 1 µm outside to 2 µm inside the reference surface, the 15 brightest intensity samples along the ray were averaged, giving a per-vertex surface intensity. A vertex was scored as enriched if this top-15 mean exceeded µ_cyto + 1.5 σ. The per-cell “surface enrichment” metric is the fraction of retained vertices scored as enriched in a given cell. A brightness ratio was also computed as the mean of the top-15 mean / µ_cyto, restricted to enriched vertices only. Since this metric focuses only on enriched vertices, it quantifies the magnitude of the SEPT6-eGFP enrichment within detected septin structures at the cell surface.

### Adherent cell imaging and stress fiber quantification

Following the same cell culture and collection protocol described above for detached cell imaging, SEPT6-eGFP MV3 cells were seeded into 35mm poly-D-lysine coated glass-bottom dish (MatTek, #P35G-1.5-14-C) at approximately 5,000 cells ml−1 based on Trypan blue counts and stored at 37 °C and 5% CO_2_ overnight to allow cells sufficient time to establish robust adhesion to the plates. The following day, medium was aspirated and replaced with either drug-treated medium or DMSO solvent-only control medium for a two-hour incubation. After incubation, cells were imaged within thirty minutes as before. Cells that were not fully in frame, made contact with other cells, appeared dead, or displayed grossly abnormal morphologies suggesting the activation of programmed cell death were all excluded from quantification. Length of the three longest stress fibers was measured per cell at the Z-stack slice image best corresponding to the basal plasma membrane (carried out with the FIJI ImageJ package). Mean pixel intensity along these stress fiber regions was measured and corrected by subtracting the mean cytoplasmic intensity at the same z-slice.

### Viability Assay

Wild-type MV3 cells were grown to 70–80% confluency in 10cm dishes, trypsinized for 3-5 minutes, and seeded into either the glass bottom 8-well µ-slides (ibidi, #80807) for adherent conditions or Bioinert 8-well µ-slides (ibidi, #80800) for detached conditions. Both slides were seeded at a density of 200 cells per well based on Trypan blue counts. Slides were incubated for 2 hours at 37 °C and 5% CO_2_, allowing adherent cells time to form attachments prior to drug treatment. Drug-treated medium or DMSO solvent-only control medium was then added to each well and incubated overnight, ensuring equal drug exposure time across adherent and detached conditions. Cells were stained with Hoeschst, added at a final concentration in the well of 3 uM (Invitrogen, #33342), and Propidium Iodide (PI), added at a final concentration of 1ug/mL (Invitrogen, #P1304MP). Live and dead cells (as identified by cellular PI signal) were counted by blinded analysts using the Cell Counter ImageJ plugin.

## Results

Given our interest in septin-mediated bleb signaling in cancer, we designed an experiment to assess the effects of known septin inhibitors on bleb signaling hubs in the context of melanoma cells. As shown in our previous work^12^, these cells do not reliably display septin hubs under normal adherent culture conditions, instead only forming them as a result of plasma membrane blebbing initiated by detachment. We previously created these conditions by embedding melanoma cells in soft bovine collagen to recreate the environment poorly-adhered amoeboid melanoma cells experience in the stroma and interstitium, imaging these samples using advanced lightsheet microscopy systems^48^. This approach, however, is limited both by the slow throughput necessitated by careful sample preparation, and the limited availability of custom-built lightsheet microscopes capable of high-resolution subcellular 3D imaging deep within collagen blocks. Moreover, such systems often require immersion of samples in large imaging chambers, significantly increasing the cost of inhibitor reagents and making matched same-day replicates impossible.

For these reasons, we opted to design experiments using a common and commercially-available spinning disk confocal microscope, imaging cells before they could establish attachment to microscopy dishes to increase throughput and reduce volumes. We treated MV3 melanoma cells expressing SEPT6-eGFP with FCF, UR214-9, REM127, or a DMSO vehicle control for 2 hours in a low-adhesion corning dish to maintain a detached state. Cells were then quickly transferred to an imaging dish and imaged. The imaging time was strictly restricted to 30 minutes to ensure that all cells experienced comparable periods of inhibitor treatment, and the microscopist was blinded to the treatment group to ensure objectivity in which cells were imaged. Blinded analysts then counted the number of detached cells clearly showing enriched septin hubs at the plasma membranes. The proportion of these “bleb signaling active” cells was calculated relative to the total number of cells imaged during that replicate (Figure S1A). While all 3 inhibitors seemed to reduce the proportion of bleb signaling active cells, only FCF produced a statistically significant difference.

Concerned that this approach was statistically underpowered, we decided to employ computer vision based quantitative methods. In our previous work, we made use of high resolution isotropic volumetric microscopy datasets to generate high fidelity 3D surface segmentations^12^. With this approach, we could then quantify the total amount of fluorescent septin signal within a given cell and calculate how much of this signal was found at the cell surface compared to the amount of signal which would be there if the total signal was equivalently distributed within the cellular volume. Unsurprisingly, we found that such an approach was not possible with our spinning disk micrographs, because their anisotropic resolution resulted in axial blurring along the top and bottom of the cell, making reliable results with this approach impossible (Figure S2). Instead, we segmented the cells and considered only the central third of their volumes – regions in which the plasma membrane was roughly orthogonal to the optical axis and thus possessed sufficient resolution to distinguish cytoplasmic signals from signals at the cell surface. For each cell, we measured signal intensity along thousands of rays that extended from the cell surface towards the cell’s centroid, normal to the cell surface. When cell surface signals along a given ray were measured to be 1.5 standard deviations above the mean cytoplasmic intensity measured by that same ray, that portion of the surface was considered to be “enriched” with septin signals. By using several thousand of these rays for each cell, we were able to fully measure the percentage of the cell’s sampled surface (i.e., the central third as explained above) that was covered by bright septin hubs. These data (Figure 1A) vastly improved the resolution of our experimental results, confirming that cells treated with FCF tended to experience considerable reductions in septin hub formation compared to DMSO controls. This approach indicated that UR214-9 was in fact significantly inhibiting septin hubs, in contrast to our human-called results. Finally, we found that REM127 showed no significant effects, producing a bimodal dataset that closely resembled that of the DMSO controls. Of note, decreasing membrane hubs did not correspond to the formation of new septin structures elsewhere in the detached cells, as the presence of cytoplasmic foci was significantly reduced by all three drugs and no ectopic filaments were seen in any treatment groups (Figure S1B). Measurement of the brightness of detected septin hubs yielded similar results to surface coverage data (Figure 1C), showing a significant reduction in FCF-treated cells, while UR214-9 produced a nonsignificant reduction, and REM127 showed no effect compared to the control. These data bring our results into better concordance with what is known of these inhibitors: UR214-9 is a derivative of FCF and thus possesses a similar capability to disrupt septin hubs, while the unrelated compound and predicted “molecular glue” REM127 does not seem to inhibit this type of higher-order septin structure.

**Figure 1:**
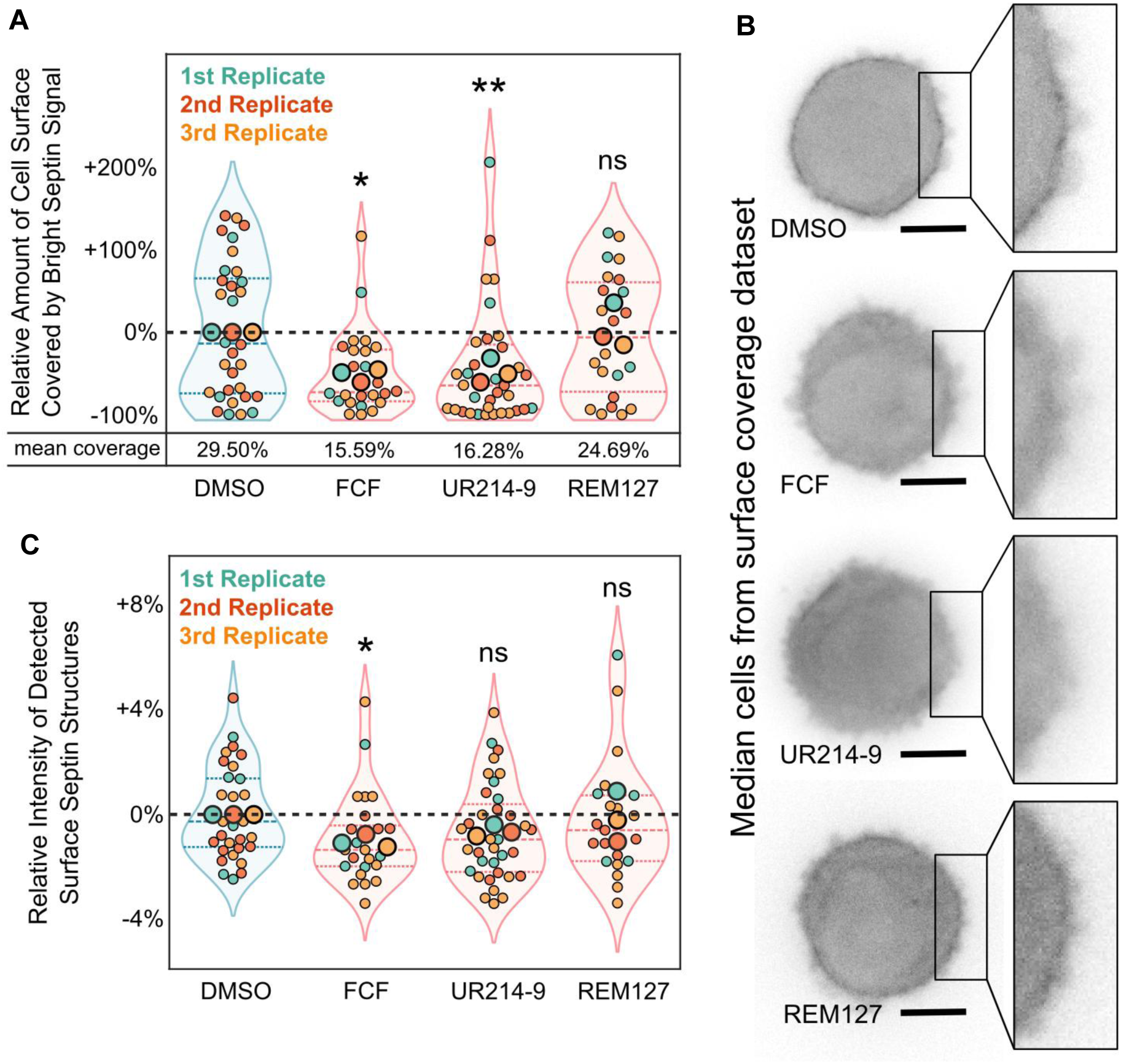
Effects of Septin Inhibitors on Bleb-Generated Septin Structures in Melanoma Cells. **a)** Effects of inhibitors on percentage of cell surface covered by bright septin structures, expressed as percent change in coverage compared to mean coverage of same-day control cells treated with DMSO alone. Small dots represent individual cells while large dots represent replicate means, with all color-coded according to replicate. Overlaid violin plots indicate probability distributions of combined datasets, with horizontal lines marking the boundaries between quartiles. Mean absolute surface coverage of all datasets shown below. Combined datasets were normality tested with Shapiro-Wilk (p=0.0084, 0.0006, <0.0001, & 0.0601). Based on these results, all three experimental groups were tested against DMSO using Mann-Whitney U tests (p=0.0271, 0.0097, & 0.7435). **b)** Median cells from all four surface enrichment datasets in figure 1a. Raw images of central z-slices are shown with no deconvolution or image enhancement beyond brightness and contrast adjustment. Scale bars represent 10 μm. **c)** Detected surface septin structures from figure 1a were measured for brightness relative to mean cytoplasmic signal. Plot shows these values expressed as percent change in brightness compared to mean brightness of detected septin structures in same-day control cells treated with DMSO alone. Graphical representations are as described for figure 1a above. Combined datasets were normality tested with Shapiro-Wilk (p=0.1199, 0.0141, 0.1394, & 0.0288). UR214-9 was variance tested against DMSO using two-tailed F-test (p=0.7419). Based on these results, FCF and REM127 were tested using Mann-Whitney U tests (p=0.0227 & 0.4621) and UR214-9 was tested using two-tailed unpaired t-test (p=0.1025). Total n=32 (DMSO), 24 (FCF), 34 (UR214-9), & 22 (REM127).

We then performed viability tests on cells treated overnight, either with one of the three inhibitors or a solvent-only control. Our previous work shows that these cells only rely upon bleb-generated septin signaling hubs for survival under detached conditions, and that septin inhibition does not meaningfully affect viability while cells are adhered^12^. The current FCF experiments replicated these previous results, with FCF treatment causing no increase in cell death when cells are grown under adherent conditions and significantly increasing death under detached conditions (Figure 2). REM127 had no effect on melanoma cell viability in either the adhered or detached state. Interestingly, UR214-9 caused significant increases in cell death in both the adhered and detached conditions, suggesting that its viability effect is not exclusively tied to septin inhibition and the disruption of the pro-survival septin hubs produced during bleb signaling, but rather a broader cytotoxic effect potentially produced by activity off-target to its effects on septins. This is in line with recent publications suggesting that UR214-9 possesses high toxicity at the dosages required for septin inhibition in both cell culture and zebrafish models^34^. Together, these results suggest that FCF is the superior inhibitor of the three currently available to researchers for inhibiting septin signaling hub formation in cancer cells undergoing bleb signaling.

**Figure 2:**
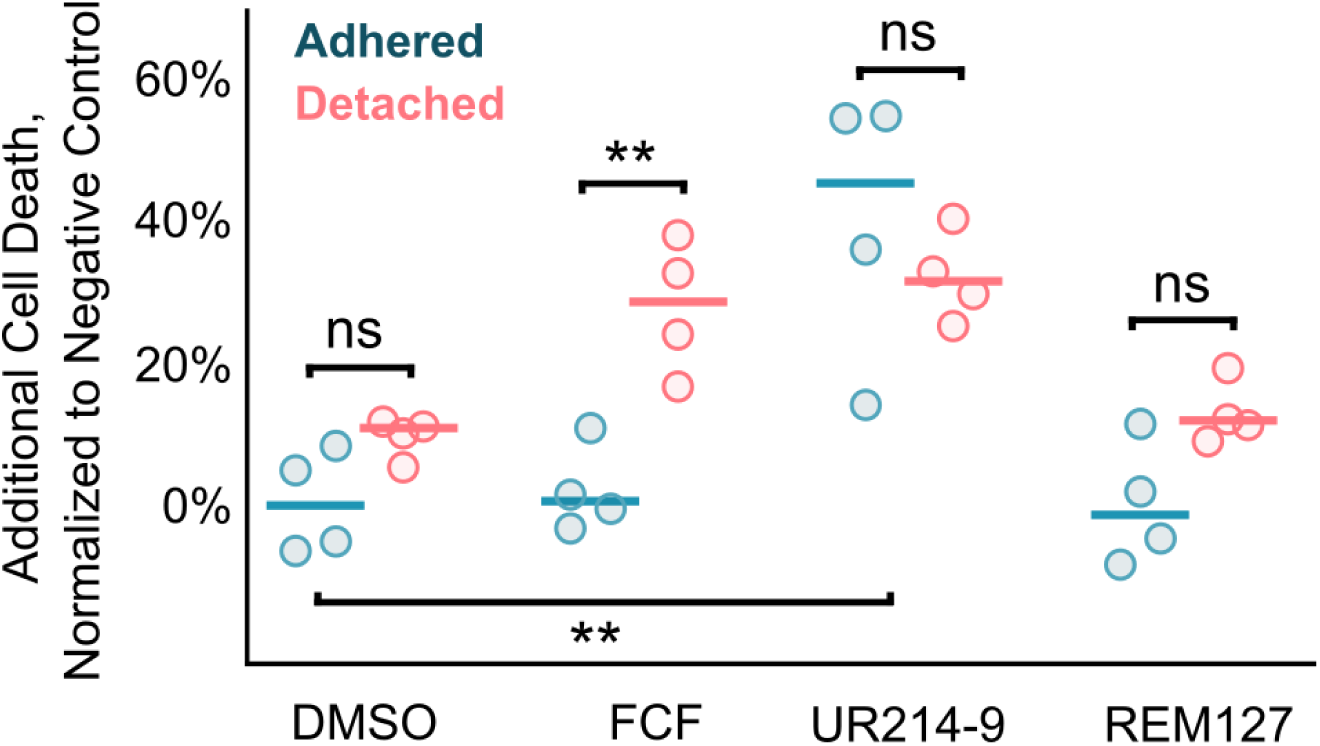
Viability Effects of Septin Inhibitors in Attached and Detached Melanoma Cells. Effects of inhibitors on cell viability in both adhered and detached states. All replicates were treated with either the indicated inhibitors or a DMSO-only negative control and grown in either adhered or detached conditions for 24 hours. Cells were then viability stained with propidium iodide and Hoechst 33342 and a blinded analyst counted and scored all propidium iodide positive cells as dead. Counts are represented here as percent dead, with replicate data normalized by subtracting the adhered negative control mean from each treatment group. All detached groups were tested against matched adhered controls using two-tailed unpaired t-tests with Welch’s correction (p=0.0818, 0.00545, 0.489, & 0.0517). Adhered UR214-9 were normality tested against adhered DMSO using Shapiro Wilk (p=0.2881 & 0.2005) and variance tested using two-tailed F-test (p=0.7124). Based on these results, it was then tested using a two-tailed unpaired t-test (p=0.0055). From left to right for all groups, total n=273, 226, 162, 184, 159, 124, 218, & 249.

In both purified in-vitro systems and living cells, FCF^29^, UR214-9^33^, and REM127^35^ have all been shown to promote septin polymerization, though the structural and biochemical mechanisms driving this change are not fully understood. In contrast to this, however, several papers have also shown that FCF treatment can instead destabilize septin polymers in some contexts^17,24,32^. To the best of our knowledge, this apparent contradiction has not been directly addressed in the literature. Because our own work on bleb signaling hubs has only ever shown FCF to be a septin destabilizer, we want to understand the contextual specificity of these opposing effects. To this end, we explored the effects of septin inhibitors on another class of higher-order septin structure known to form in MV3 melanoma cells: the septin filaments decorating stress fibers.

To induce stress fiber formation, the same MV3 line was seeded in imaging dishes overnight to allow sufficient time to adhere and form structures. The following day cells were subjected to the same drug treatment and imaging protocol as in Figure 1 above. The resulting images were analyzed using ImageJ macros to record the intensity and length of septin decorated fibers. Interestingly, our results showed that all three drugs caused clear elongation of septin filaments compared to DMSO controls (Figure 3A). Septin intensity along the fiber was modestly reduced by all drug treatments, though not to a statistically significant degree. Together, these results suggest an increase in end-on-end septin polymerization, without an accompanying increase in lateral association between septin rods or filaments. This strongly points to the duality of septin inhibitor effects not being exclusively dependent upon the cell type being treated, but instead upon the particular septin structure being considered. Moreover, this highlights the importance of considering the distinct classes of higher-order septin structures within cells and the disparate possible effects of septin targeting drugs on such structures.

**Figure 3:**
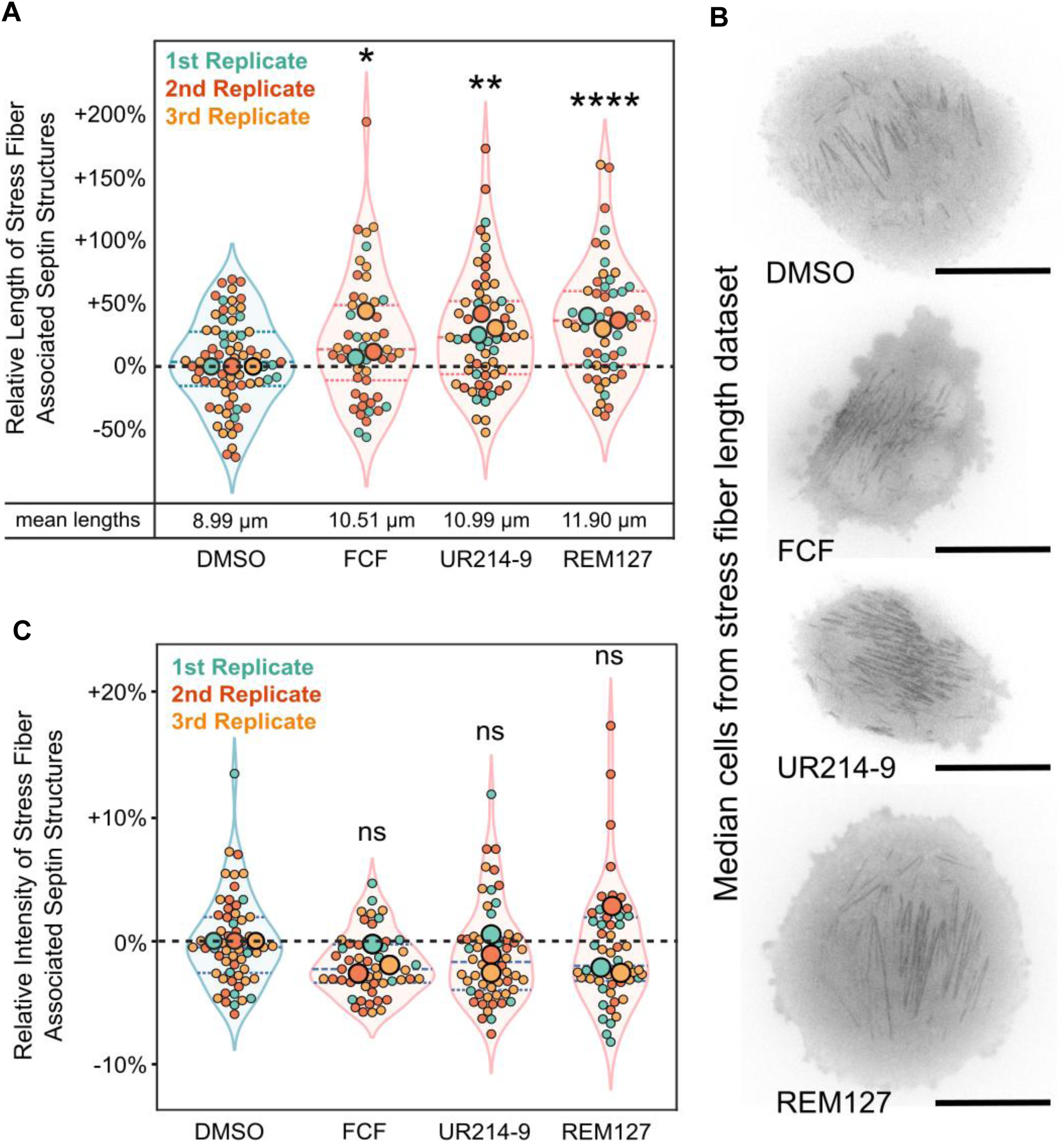
Effects of Septin Inhibitors on Stress Fiber Associated Septin Filaments in Melanoma Cells. **a)** Effects of inhibitors on lengths of septin filaments decorating stress fibers at the basal membrane of adhered melanoma cells, expressed as percent change in length compared to mean length of same-day control cells treated with DMSO alone. Small dots represent individual cells while large dots represent replicate means, with all color-coded according to replicate. Overlaid violin plots indicate probability distributions of combined datasets, with horizontal lines marking the boundaries between quartiles. Mean absolute lengths from all datasets shown below. Combined datasets were normality tested with Shapiro-Wilk (p=0.4559, 0.0155, 0.0410, & 0.0755) and REM127 was variance tested against DMSO using a two-tailed F-test (p=0.0399). Based on these results, FCF and UR214-9 were tested against DMSO using Mann-Whitney U tests (p=0.0295, & 0.0032) and REM127 was tested using a two-tailed unpaired t-test with Welch’s correction (p<0.0001). **b)** Median cells from all four surface enrichment datasets in figure 3a. Raw images of central z-slices are shown with no deconvolution or image enhancement beyond brightness and contrast adjustment. Scale bars represent 10 μm. **c)** Septin filaments from figure 3a were measured for brightness relative to mean cytoplasmic signal in the same cell and expressed as percent change from mean filament brightness of same-day control cells treated with DMSO alone. Graphical representations are as described for figure 3a above. Combined datasets were normality tested with Shapiro-Wilk (p=<0.0001, <0.0001, <0.0001, & <0.0001). Based on these results, FCF, UR214-9, and REM127 were tested using Mann-Whitney U tests (p=0.1061, 0.2343 & 0.3962). Total n=69 (DMSO), 50(FCF), 58(UR214-9), & 52(REM127).

## Discussion

Septin dynamics is a field of increased interest due to the many cellular processes implicated, but the complexity of septin oligomer formation and higher-order polymerization makes these processes quite difficult to elucidate. Our study provides insights into the effects of septin inhibitors on higher-order structures in different cellular contexts in cancer cells. Prior studies using living cells and purified septin oligomers exposed to FCF, REM127 and UR214-9 *in vitro* have demonstrated that these drugs can polymerize and stabilize septin structures ^33,35,49^. Counterintuitively, the opposite effect has also been shown, with FCF treatment causing the breakdown of higher-order septin structures in cells^12,17,24,32^. Here we show that in melanoma cells, septin targeting drug treatment results in both context-dependent effects, promoting polymerization in some cases while inducing destabilization in others. This highlights that the disparity does not seem to be due to different cell types, but instead to the drugs having alternative effects on different classes of septin structures. What characteristics distinguish these classes and by what mechanism these disparate effects are achieved are both open questions.

One possibility is that the septin proteins making up these different classes are distinct from one another and thus have different sensitivities to these drugs. This seems reasonable at first glance because of the wide variety of different septin proteins produced by mammalian cells. Indeed, there are 13 different septin-coding genes in the human genome, many of which have over a dozen different splice variants. This complexity results in hundreds of different possible species of septin oligomer, even if only considering canonical hexameric and octameric complexes. It is possible that different oligomer species might make up the septin structures at bleb signaling hubs and stress fibers, resulting in the observed differences in drug sensitivities.

A key observation arguing against this idea, however, is that all three inhibitors produced remarkably similar phenotypes despite having distinct predicted binding modes. One might have expected these opposing effects to be a specific quirk of FCF, but here we show that UR214-9 and REM127 share this duality as well. In every case, drug treatment increased filamentous septin structures at stress fibers while failing to enhance bleb signaling hub formation. This convergence is striking because the inhibitors are thought to interact with septins through entirely different mechanisms. FCF is a substituted phenylurea compound predicted to interact with the GTP-binding pockets of SEPT2, SEPT3, and SEPT7^31^. Conversely, REM127 is a thiadiazole derivative shown to bind specifically between the NC interface of SEPT6 and SEPT7^35^, an entirely different region of the oligomeric complex. For differential oligomer composition to explain the observed phenotype, hub-associated oligomers would need to be simultaneously resistant to the polymerization-promoting effects of multiple mechanistically-distinct drugs. While this possibility cannot be excluded, it appears less parsimonious than other explanations.

Alternatively, rather than considering the nature of the oligomers being assembled into higher-order structures, one might instead consider the nature of the higher-order interactions between these oligomers. As already discussed, all three compounds have demonstrated the ability to polymerize septins end-on-end *in vitro*, a behavior they also promote at stress fibers in live cells. Importantly, the formation of septin structures at stress fibers has been conclusively demonstrated to require end-on-end polymerization^7^. But perhaps the higher-order structures seen at bleb signaling hubs do not rely on this classic polymerization pattern. Indeed, stabilized end-on-end polymerization might make such higher-order structures less likely to occur. For instance, weak, multivalent interactions between c-terminal extensions or other domains might drive hub formation after oligomers are recruited to the membrane via bleb-generated plasma membrane curvature. Supporting this possibility are several recent papers suggesting septin structures might undergo liquid-liquid phase separation in this way, specifically at adherens junctions^50,51^. This is quite interesting, as all previous papers we were able to identify in which FCF treatment resulted in higher-order structure disruption, dissipating bright septin signals to produce a more uniform cytoplasmic distribution, have been specifically studying septins at adherens junctions. If hubs and adherens junctions are similarly destabilized by increased end-on-end polymerization, it suggests they might both be made up of phase-separated septin condensates. This suggests many new hypotheses regarding the higher-order structure of bleb-generated septin signaling hubs and the mechanisms by which they scaffold and regulate signaling.

Beyond such speculation, this work shows that of the three inhibitors considered in this study, FCF is superior in the context of inhibiting bleb signaling in detached cancer cells. UR214-9, while producing significant inhibition of hub formation, results in off-target toxicity. REM127 does not share this cytotoxicity but also seems to lack the ability to strongly inhibit hubs or produce the resulting decrease in viability upon detachment. A broader dose response study would strengthen these conclusions, though it should be noted that UR214-9 dosage in this study yielded modestly weaker septin hub inhibition. This means that further dose reduction in the hopes of avoiding toxicity would produce even less on-target effects relative to FCF. Similarly, the REM127 dose was sufficiently high to produce the strongest pro-polymerization effects of all three drugs in adhered cells, suggesting the lack of effect on hubs was not due to dosage being below a threshold necessary for on-target septin effects. Titrating dosage higher to see if this pattern holds true would nonetheless be interesting, as would studying the other compounds in the Res19-T and UR214 series to determine if they can efficiently disrupt hubs. Regardless, as our recent work suggests that septin inhibition might be a clinically effective means of targeting metastatic cancer cells employing septin-mediated bleb signaling, the current study highlights a continued need for improved septin inhibitors that can effectively and specifically target the distinct classes of septin higher-order structures responsible for this disease-related phenotype.

Septins structures were first visualized in *Saccharomyces cerevisiae* forming filamentous rings around the cleavage furrow during cytokinesis^52^. Septins are now known to form a variety of structures with diverse functions in animal cells, acting as diffusion barriers in primary cilia^21,53^, coordinating syncytial blastoderm organization in *Drosophila*^54^, regulating calcium release in neurons^35^, contributing to myelination in oligodendrocytes^55^, modulating mitochondrial fission^56^, initiating platelet activation^57^, regulating midbody-based cell fate specification^58^, and maintaining barrier integrity and cell polarity via adherens junctions^51,59^. Given this broad functional distribution across organisms, it is worth considering not only the cellular contexts and roles of these septin assemblies, but more specifically which of the putative structural classes described here they might fall within – not only to better design pharmacological modifications of such functions, but to better understand their underlying molecular biology.

## Acknowledgements

The authors would like to thank Dan Dickinson for the generous use of his imaging facilities.

---

**Supplemental Figure S1:**
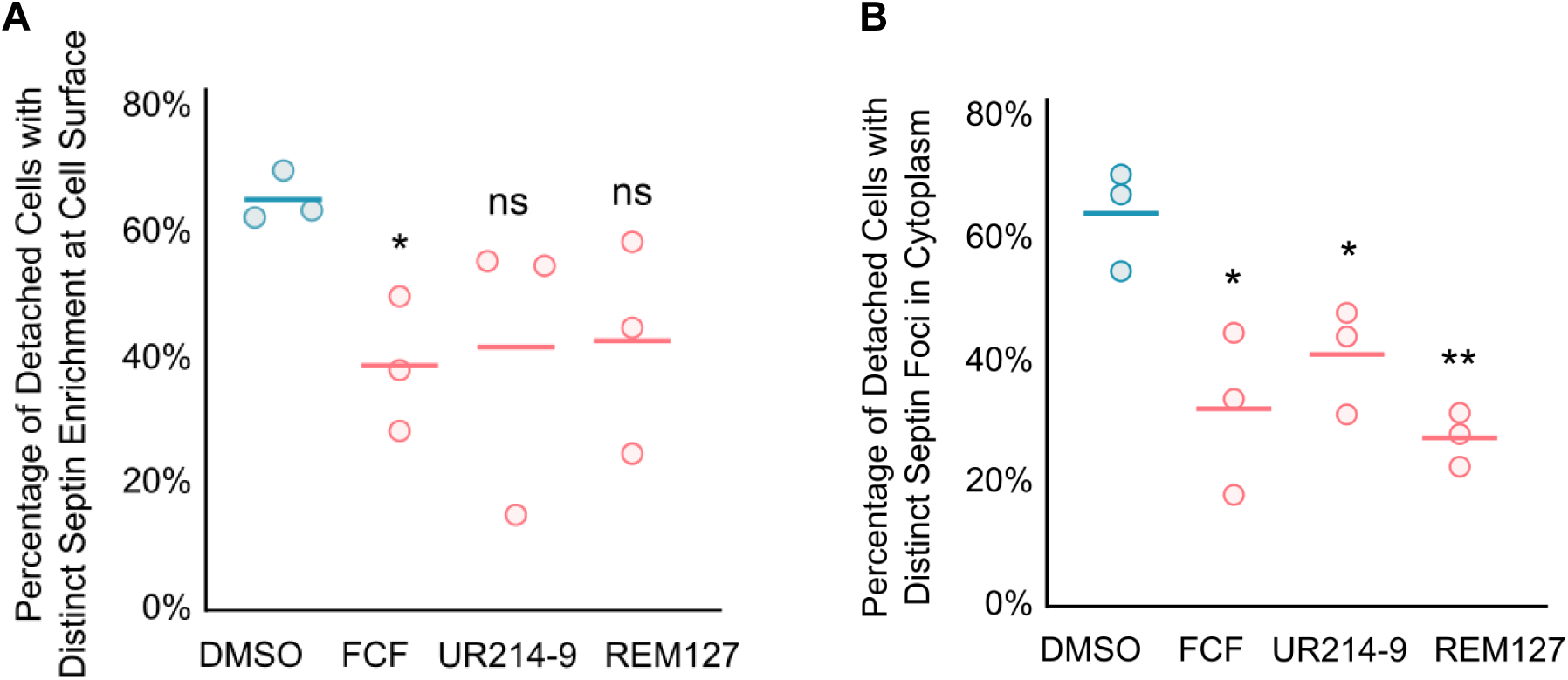
Human Calls of Septin Localization in Detached Melanoma Cells. **a)** Percentage of cells across treatment groups showing distinct enrichment of SEPT6-eGFP signal at the surface, as determined by blinded human analysts. Cells across all groups were imaged in matched, same-day replicates on 3 separate days. Cells were detached for two hours in the presence of either drugs or in DMSO-only control medium, after which they were immediately imaged in strict 30 minute windows to ensure no difference in treatment time between groups. All groups were normality tested with Shapiro-Wilk (p= 0.2693, 0.8920, 0.0297, & 0.7916) and variance tested against control with two-tailed F-test (p=0.2485, 0.0595, & 0.1081). Based on these results, FCF and REM127 were then tested with two-tailed unpaired t-test with Welch’s correction (p=0.0376, & 0.1411) and UR214-9 with two-tailed Mann-Whitney (p=0.100). b) Percentage of the cells from figure **S1a** that showed distinct foci of SEPT6-eGFP signal within the cytoplasm, as determined by blinded human analysts. All groups were normality tested with Shapiro-Wilk (p= 0.3839, 0.7966, 0.4316, & 0.7615) and variance tested against control with two-tailed F-test (p=0.5645, 0.9608, & 0.4412). Based on these results, all 3 experimental groups were tested against DMSO with two-tailed unpaired t-tests (p=0.0245, 0.0294, & 0.0026). Total n= 55 (DMSO), 54 (FCF), 95 (UR214-9), & 66 (REM127)

**Supplemental Figure S2:**
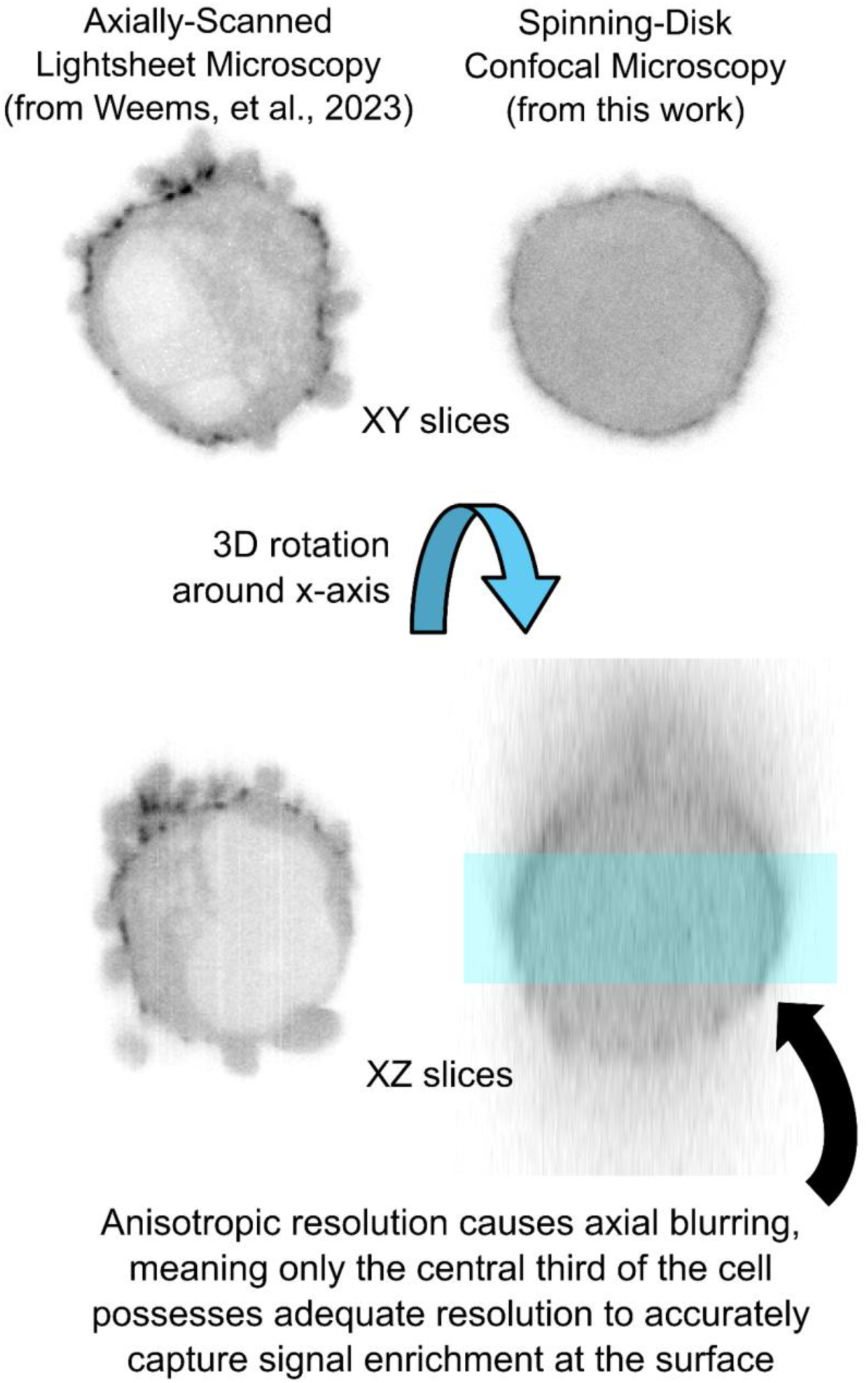
Visual Representation of Z-Resolution in ASLM vs Spinning-Disk.

